# Disruption of perineuronal nets increases the frequency of sharp wave ripples

**DOI:** 10.1101/124677

**Authors:** ZhiYong Sun, P. Lorenzo Bozzelli, Adam Caccavano, Megan Allen, Jason Balmuth, Stefano Vicini, Jian-Young Wu, Katherine Conant

## Abstract

Hippocampal sharp wave ripples (SWRs) represent irregularly occurring synchronous neuronal population events that are observed during phases of rest and slow wave sleep. SWR activity that follows learning involves sequential replay of training-associated neuronal assemblies and is critical for systems level memory consolidation. SWRs are initiated by CA2 or CA3 pyramidal cells and require initial excitation of CA1 pyramidal cells as well as subsequent participation of parvalbumin (PV) expressing fast spiking (FS) inhibitory interneurons. These interneurons are relatively unique in that they represent the major neuronal cell type known to be surrounded by perineuronal nets (PNNs), lattice like structures composed of a hyaluronin backbone that surround the cell soma and proximal dendrites. Though the function of the PNN is not completely understood, previous studies suggest it may serve to localize glutamatergic input to synaptic contacts and thus influence the activity of ensheathed cells. Noting that FS PV interneurons impact the activity of pyramidal cells thought to initiate SWRs, and that their activity is critical to ripple expression, we examine the effects of PNN integrity on SWR activity in the hippocampus. Extracellular recordings from the stratum radiatum of 490 micron horizontal murine hippocampal hemisections demonstrate SWRs that occur spontaneously in CA1. As compared to vehicle, pretreatment (120 min) of paired hemislices with hyaluronidase, which cleaves the hyaluronin backbone of the PNN, decreases PNN integrity and increases SWR frequency. Pretreatment with chondroitinase, which cleaves PNN side chains, also increases SWR frequency. Together, these data contribute to an emerging appreciation of extracellular matrix as a regulator of neuronal plasticity and suggest that one function of mature perineuronal nets could be to modulate the frequency of SWR events.

## Introduction

Hippocampal sharp wave ripples (SWRs) represent synchronous neuronal population events characterized by a low frequency large amplitude deflection and an associated high frequency ripple oscillation (Buzsaki, 1986). SWRs have been implicated in the rapid sequential replay of neuronal assemblies that were initially activated during a learning experience (Alme et al., 2014; Colgin, 2016; O’Keefe and Dostrovsky, 1971; Wilson and McNaughton, 1994). During these events, replay of assembly sequences occurs at an accelerated rate and is thought to be critical for memory consolidation and transfer (Skaggs et al., 1996). Consistent with this are studies showing that disruption of SWR activity during sleep impairs memory for previously experienced information and that sustained increases in SWR activity are observed during slow wave sleep that follows learning (Ego-Stengel and Wilson, 2010; Eschenko et al., 2008; Girardeau et al., 2009). In addition, sequential replay reverberates between varied brain regions implicated in the acquisition and/or longterm storage including hippocampus and cortical projection areas (Ji and Wilson, 2007; Place et al., 2016; Rothschild et al., 2017). Intriguingly, sequential replay of previously activated neuronal assemblies occurs in a bidirectional manner, and forward replay may facilitate the ability to plan for future events (Dragoi and Tonegawa, 2011; Sadowski et al., 2011). Proper timing of SWR sequences likely impacts information transfer or planning and is thus an area of study within the field.

Pyramidal cells (PCs) of CA3 have been implicated in the initiation of hippocampal SWRs both *in vivo* and *in vitro* (Buzsaki, 2015). The sharp wave is generally transmitted from CA3 to CA1 and represents an excitatory event, while the ripple is generated locally in CA1 by ripple –frequency activity of PV expressing fast spiking interneurons (Schlingloff et al., 2014; Whittington et al., 1995). SWRs are initiated when a sufficiently large number of PCs is activated within a narrow time window, and enhancement of cellular excitability through modification of ACSF can increase SWR rate (Schlingloff et al., 2014).

PV expressing basket cells not only generate ripple frequency oscillations, but influence the excitability of principal cells that initiate sharp wave ripples. In one study that examined pyramidal cell excitability in mice engineered so that AMPA mediated excitation of PV interneurons was selectively suppressed, it was observed that pyramidal cells were more likely to fire (Racz et al., 2009). PV interneurons also represent the class of hippocampal interneurons that have been implicated in ripple expression (Schlingloff et al., 2014). This was demonstrated by experiments with perisomatic injection of ω-agatoxin, which targets voltage dependent calcium channels on PV but not cholecystokinin basket cell axon terminals. Ripples disappeared when ω-agatoxin was injected into the stratum pyramidale (Schlingloff et al., 2014).

PV interneurons represent the major population of neurons that is ensheathed in a perineuronal net, a specialized extracellular matrix comprised of a hyaluronin backbone that surrounds the soma and proximal dendritic area (reviewed in (Kwok et al., 2011; Pantazopoulos and Berretta, 2016; Sorg et al., 2016)). The presence of the PNN may enhance PV interneuron excitability (Balmer, 2016) as well as the frequency of inhibitory currents in pyramidal neurons (Slaker et al., 2015). Though the mechanism(s) by which this is achieved are not completely understood, PNNs may serve to prevent diffusion of glutamate released at PV post-synaptic contacts and/or restrict the lateral mobility of GluAs on this population (Balmer, 2016; Bikbaev et al., 2015; Frischknecht et al., 2009; Slaker et al., 2015). Consistent with the latter possibility, the diffusion constant of GluA1 drops as the subunit reaches hyaluronin-enriched area. Hyaluronidase digestion increased both the diffusion coefficient and the total surface area covered by individual subunits (Frischknecht et al., 2009).

PNN integrity varies as a function of normal physiology and pathology (Kwok et al., 2011; Pantazopoulos and Berretta, 2016). Developmental maturation of the PNN correlates with closure of critical periods of plasticity in which appropriate experience molds neuronal network organization (Pizzorusso et al., 2002). PNN formation is neuronal activity dependent (Dityatev et al., 2007), and region specific development is influenced by relevant neuronal activity or lack thereof. PNNs are more numerous in song bird species with closed end song learning as opposed to those that display extensive adult vocal plasticity (Cornez et al., 2017). Moreover, PNN deposition and critical period closure in the visual cortex are both reduced by dark rearing (Gianfranceschi et al., 2003; Hockfield et al., 1990; Mower, 1991), and effects of monocular deprivation are mimicked by matrix degradation (Oray et al., 2004). In addition, in the mouse barrel cortex, whisker trimming during the first post-natal month is associated with a reduction in aggrecan rich PNNs (McRae et al., 2007). Of interest, the monoamine reuptake inhibitor fluoxetine has been shown to reduce PNN integrity and to reopen the critical window for visual plasticity (Guirado et al., 2014; Maya Vetencourt et al., 2008). Altered PNN integrity is also observed in select psychiatric conditions, including depression (Pantazopoulos et al., 2015) and may also occur following seizure, trauma or infection (Franklin et al., 2008; Hsieh et al., 2016; Pollock et al., 2014). Consistent with these observations, levels of PNN degrading enzymes may be increased in the background of injury or seizure activity (Mayer et al., 2005; Pollock et al., 2014).

Despite the likely importance of the PNN to PV activity, and the role of PV interneurons in SWR expression, few if any studies have explicitly examined effects of PNN integrity on SWR events. The present study addresses the impact of hyaluronidase processing of the PNN on SWR frequency. We use acute hippocampal slices of thickness sufficient for the expression of spontaneous events (Schlingloff et al., 2014), and paired dorsal to ventral hemislices that are concomitantly pre-incubated in control or hyaluronidase-containing ACSF.

## Materials and Methods

### Slice preparation

6-8 week old male and female C57/Bl6 mice were obtained from Jackson laboratories and used to prepare paired hippocampal hemi-slices in accordance with National Institutes of Health guidelines and a protocol that had been approved by the Institutional Animal Care and Use Committee at Georgetown University Medical Center. Following deep isoflourane anaesthesia, animals were rapidly decapitated. The whole brain was subsequently removed and chilled in cold (0° C) sucrose-based cutting artificial cerebrospinal fluid (sACSF) containing (in mM) 252 sucrose; 3 KCL; 2 CaCl2; 2 MgSO_4_; 1.25 NaH2PO4; 26 NaHCO3; 10 dextrose and bubbled by 95% O_2_, 5% CO_2_. Hippocampal slices (490 um thick) were cut in horizontal sections from dorsal to ventral brain with a vibratome (Leica, VT1000S). Horizontal sections were then bisected to obtain paired hemislices for subsequent incubation of one versus the other paired hemislice with control or enzyme containing ACSF. ACSF contained (in mM) NaCl, 132; KCl, 3; CaCl2, 2; MgSO4, 2; NaH2PO4, 1.25; NaHCO3, 26; dextrose 10; and saturated with 95% O2, 5% CO2 at 26° C. Slices were incubated for at least 120 minutes before being moved to the recording chamber.

### Chemicals and reagents

Hyaluronidase and chondroitinase ABC were purchased from Sigma Chemical (St. Louis, MO; catalogue numbers H4272 and C3667). Lyophilized preparations were reconstituted in ACSF just prior to slice incubation. The final concentration of hyaluronidase in the incubating solution was .25 gm/ml and the final concentration of chondroitinaseABC was 0.1U/ml. Both concentrations were based on previously published *in vitro* studies (Filous et al., 2014) (Bikbaev et al., 2015).

### PV and PNN staining

125 μm horizontal brain sections were cut using a vibratome under the same conditions as those mentioned above. Following a 2-hour treatment with hyaluronidase or vehicle containing carbogenated ACSF, free-floating sections were fixed overnight in 4∼ PFA at 4°C. Sections were then washed twice with 1X-PBS and permeabilized with 1X-PBS containing .5% Triton-X overnight at 4°C. Sections were then blocked with 1X-PBS containing 20% goat serum and .5% Triton-X overnight at 4° C. Sections were incubated with mouse anti-parvalbumin primary antibody (1:300, Sigma-Aldrich, P3088) in 1X-PBS containing 5% goat serum overnight at 4°C. Following three washes with 1X-PBS, sections were incubated with goat anti-mouse IgG Alexa Fluor 555 (1:750, ThermoFisher Scientific, A21425) and fluorescein-labeled *Wisteria floribunda* lectin (1:200, Vector Laboratories, FL-1351) in 1X-PBS containing 5% goat serum for 2-hours at room temperature. Sections were then washed three times with 1X-PBS, dried, and Hydromount (National Diagnostics, HS-106) was applied followed by a coverslip and allowed to dry several days at 4°C prior to confocal imaging.

### Confocal Microscopy

Tissue was imaged according to a standardized protocol (Slaker et al., 2016b). Images were acquired using a Leica SP8 laser scanning confocal microscope with an oil immersion, 20X objective and a 0.75 numerical aperture. Leica Application Suite was used for image acquisition. Laser intensity, gain, and pinhole settings were kept constant for all samples. Images were taken through a z-plane (8.5 μm) containing 20 stacks (0.4 μm/stack). Images were transferred to ImageJ (NIH) software and representative images were obtained using the ZProject maximum intensity projection.

### Local field potential (LFP) recordings

Low resistance glass microelectrodes were used for LFP recordings of the SW/R signals. The electrodes were pulled with a Sutter P87 puller with 6 controlled pulls, resulting in an approximate 150K tip resistance. Electrodes were filled with 1M NaCl in 1% agar, which prevents leakage of the electrode solution that could potentially alter conditions at the recording site. The recordings were done in a submerged chamber, and slices were perfused on both sides at a high flow rate (20 ml /min). In our recording arrangement SWR signals have an adequate signal-to-noise ratio. Hippocampal slices from dorsal and ventral locations demonstrated SWR activity. Comparisons between control and treated slices were performed with matched left and right slices of specific dorsal to ventral level sections. Thus, recordings were performed using control and treated pairs prepared at the same time and with the same cutting or recording ACSF.

### Data analysis

SWR frequency was quantified through complementary approaches. 1) Raw LFP traces were filtered between 0.5-30 Hz, and the SWR peaks were counted manually by a hypothesis-blind investigator. 2) SWR events were also characterized via a custom MATLAB algorithm based on previously published techniques (Csicsvari et al., 1999; Eschenko et al., 2008; Siapas and Wilson, 1998). A Gaussian FIR bandpass filter with corrected phase delay was applied to the LFP in two frequency bands: 2-30Hz and 80-250Hz to extract the low-frequency (LF) SW and high-frequency (HF) ripple components, respectively. The 80Hz lower cutoff for the ripple band was chosen to better match the room temperature recordings (Papatheodoropoulos et al., 2007). For both the LF and HF filtered signals, the root mean square (RMS) was computed every 5ms in a 10ms moving window. Based on published studies (Ccedil;aliskan et al., 2015; Maier et al., 2003), the threshold for peak detection was 3-4 SDs of the RMS signal as indicated. Peak duration was defined as the contiguous period around a peak where the RMS signal remained 2 SDs above baseline. In this way, both SW and ripple events could be individually detected. SWR events were defined as a subset of these, requiring both a simultaneous SW and ripple event. The duration of SWR events was defined as the greater of the overlap of simultaneous SW and ripple events. Event power was calculated by integrating either the LF, HF, or unfiltered LFP for the SW, ripple and SWR events, respectively.

### Statistical analysis

Data was entered into a Graph Pad Prism program and statistical analysis performed using Student’s t test for 2 group comparisons or ANOVA for comparisons of more than 2 groups. Significance was set at p < 0.05.

## Results

### I. Spontaneous SWR activity in 490 μm hippocampal slices

Spontaneous SWR activity has previously been reported in *ex vivo* hippocampal slices of thickness thought sufficient for adequate network complexity in elegant work by Schlingloff and colleagues (Schlingloff et al., 2014). As shown in figure 1, we also observe spontaneous SWR activity in relatively thick (490 μm) sections. Raw SWR signals from two electrodes that were placed within the proximal section of the CA1 stratum radiatum are shown. As can be appreciated from this 15 second epoch shown, events with varied inter-event intervals and morphology are apparent at both recording sites. Twenty six events are displayed with nine expanded to show the associated ripple component.

**Figure 1.**
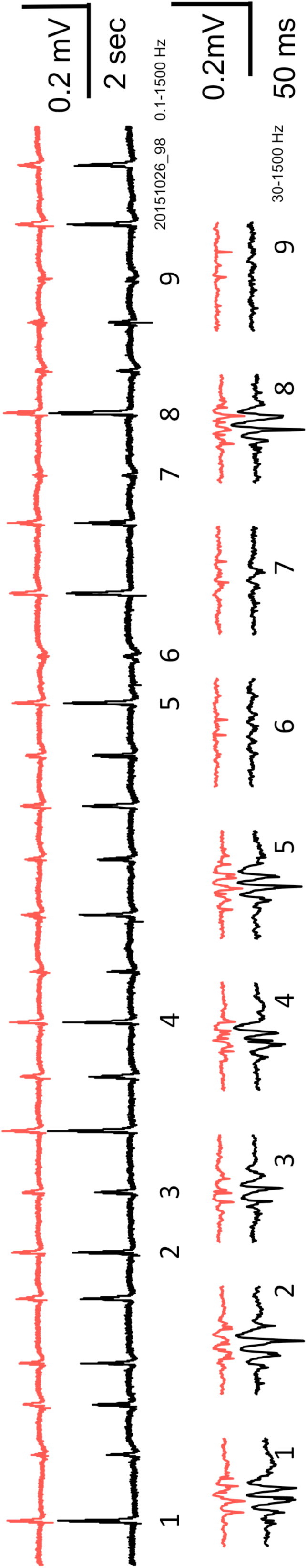
Spontaneous SWRs in 490 mM horizontal hippocampal slices. *Top:* raw LFP traces (red and black) showing sharp waves from two recording locations in CA1 (filtered; 0.1 to 1500 Hz). *Bottom:* examples of ripples from the top trace are displayed on an expanded time scale (filtered; 30 to 1500 Hz). Note that in this 15 second section of recordings there are 26 SWRs, some large events (note 1,4,8) and some small events (note 6,7,9).

**Figure 2.**
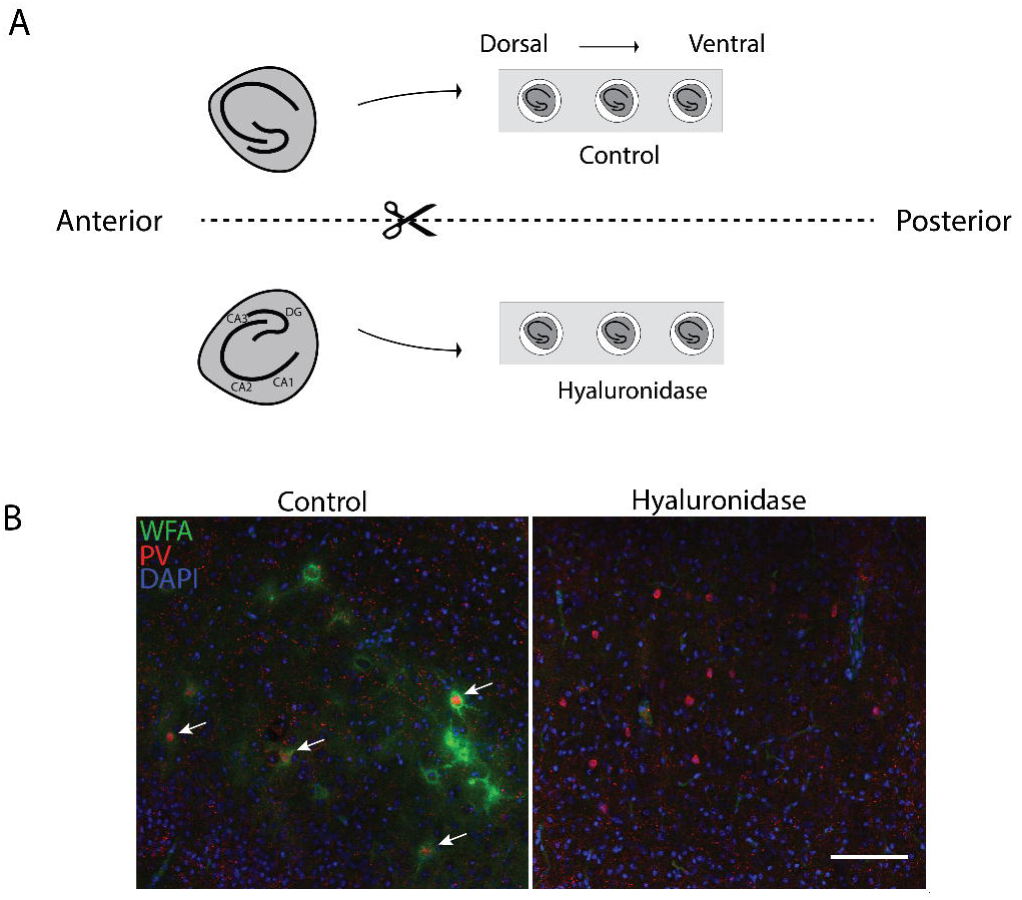
Horizontal slice preparation and staining. Horizontal slices were prepared and bisected, with matched hemisections placed in one of two recovery solutions (ACSF or ACSF with hyaluronidase). Following a 2h recovery period, hemislices were used for additional studies. A schematic of the set up is shown in 2A, and representative staining of control and hyaluronidase treated slices is shown in 2B. PNN staining is shown in the green channel and PV in the red. A qualitative loss of PNN signal surrounding PV positive neurons (white arrows) can be appreciated following hyaluronidase treatment. The scale bar represents 100 μm.

**Figure 3.**
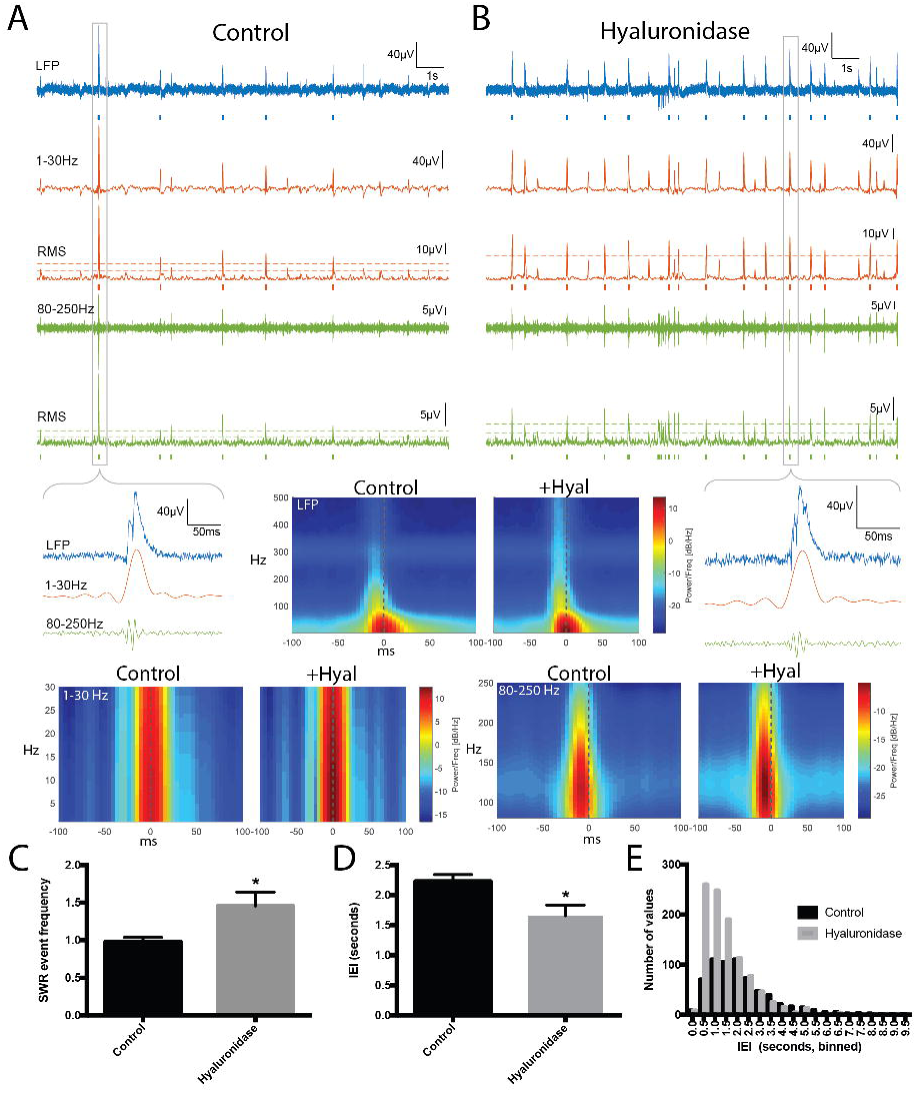
SWR event frequency in hyaluronidase treated slices. A: 1^st^ trace: Control slice 15s recording of LFP (filtered 0.5-1500Hz). Blue bars indicate detected SWR event, defined as a coincident SW and ripple event. 2^nd^ trace: lower frequency SW component filtered 1-30Hz. 3^rd^ trace: Root mean square of SW filtered from 1-30Hz. Dashed lines indicate 2 and 4 SD above baseline for event detection. Red bars below indicate detected SW event. 4^th^ trace: high frequency ripple component filtered from 80-250Hz. 5^th^ trace: Root mean square of ripple filtered 80-250Hz. Dashed lines indicate 2 and 4 SD above baseline for event detection. Green bars below indicate detected ripple event. B shows the same features for Hyaluronidase treatment. Zoomed in views of indicated SWR events are shown, as are spectrogram averages for the indicated frequency ranges in all detected SWR events from 8 control and 8 hyaluronidase pretreated slices (time-locked to the SW peak; n= 743 control events and 1114 hyaluronidase events). C shows the fold change in average SWR event frequency for two to three paired slices from each of 3 different animals (n=8 slices per group), and overall SWR inter-event interval results from the 8 control and 8 treated slices is shown in D. The difference between control and treatment SWR frequency in 3C and D is significant (*p* < 0.05). A frequency histogram of inter-event intervals is shown in E.

### II. PNN staining in hippocampal slices

The ability of hyaluronidase to disrupt PNN integrity has been demonstrated in varied settings. Due to the potential for reduced enzyme activity in the presence of carbogenated ACSF, however, we examined PNN integrity in slices that were incubated for 2h in control or hyaluronidase containing carbogenated ACSF. The experimental set up (2A) involved two separate chambers containing multiple wells so that matched horizontal hemisection pairs could recover in one of the two solutions. This set up would be maintained for future studies of SWR activity following the recovery period. In these experiments, 125 μm sections were incubated for 2h with (0.25 grams/ml) hyaluronidase, a concentration that is consistent with that recently used in the treatment of hippocampal cultures (Bikbaev et al., 2015), then formalin fixed and then stained to detect PNN ensheathed PV neurons. Shown in 2B are representative immunostaining results from control and hyaluronidase pretreated conditions. White arrows indicate PV positive neurons surrounded by PNN in control sections. PNN signal is not appreciable following 2h treatment with hyaluronidase.

### III. SWR ripple event frequency is increased following exposure to hyaluronidase

We next examined the frequency of SWR events in paired control and hyaluronidase treated hemislices. Representative traces are shown in 3A and B, with SW events in red and ripples in green. Events are underlined by tics for SWs and ripples that were detected with a 4 SD threshold. Tic length reflects event duration. A zoomed in view is shown for one event, and a spectrographic representation corresponding to the average of all detected events across 15 15s epochs for 8 control and 8 treated slices is also shown as indicated (Control or +Hyal). In 3C, we show average SWR event frequency results from 8 control and 8 hyaluronidase pretreated slices. This endpoint is significantly increased in the hyaluronidase group (*p* < 0.05). In 3D we show average inter-event interval (IEI) data as an additional form of this measure and in 3E we show a frequency distribution histogram comparing IEIs in control and hyaluronidase treatments. An increase in relatively short IEIs can be appreciated in the hyaluronidase pretreated slices.

### IV. SWR frequency is also increased following exposure to chondroitinase

Multiple *in vitro* and *in vivo* studies have also utilized chondrotinase, which targets chondroitin sulfate glycosaminoglycans, to reduce or eliminate PNNs (Lin et al., 2008). Additional studies were therefore performed with chondroitinase to further support a connection between PNN integrity and SWR event frequency. Results are shown in figure 4. Representative traces are shown in 4A-B, and in 4C the average SWR event frequency in 4 control and 4 chondroitinase slices is shown. Based on visual inspection (Koniaris et al., 2011) and consistency with hand count results, the detection threshold for data shown from these experiments was based on SD=3. For both SD3 and SD4, the difference between the groups is significant at *p* < 0.05.

**Figure 4.**
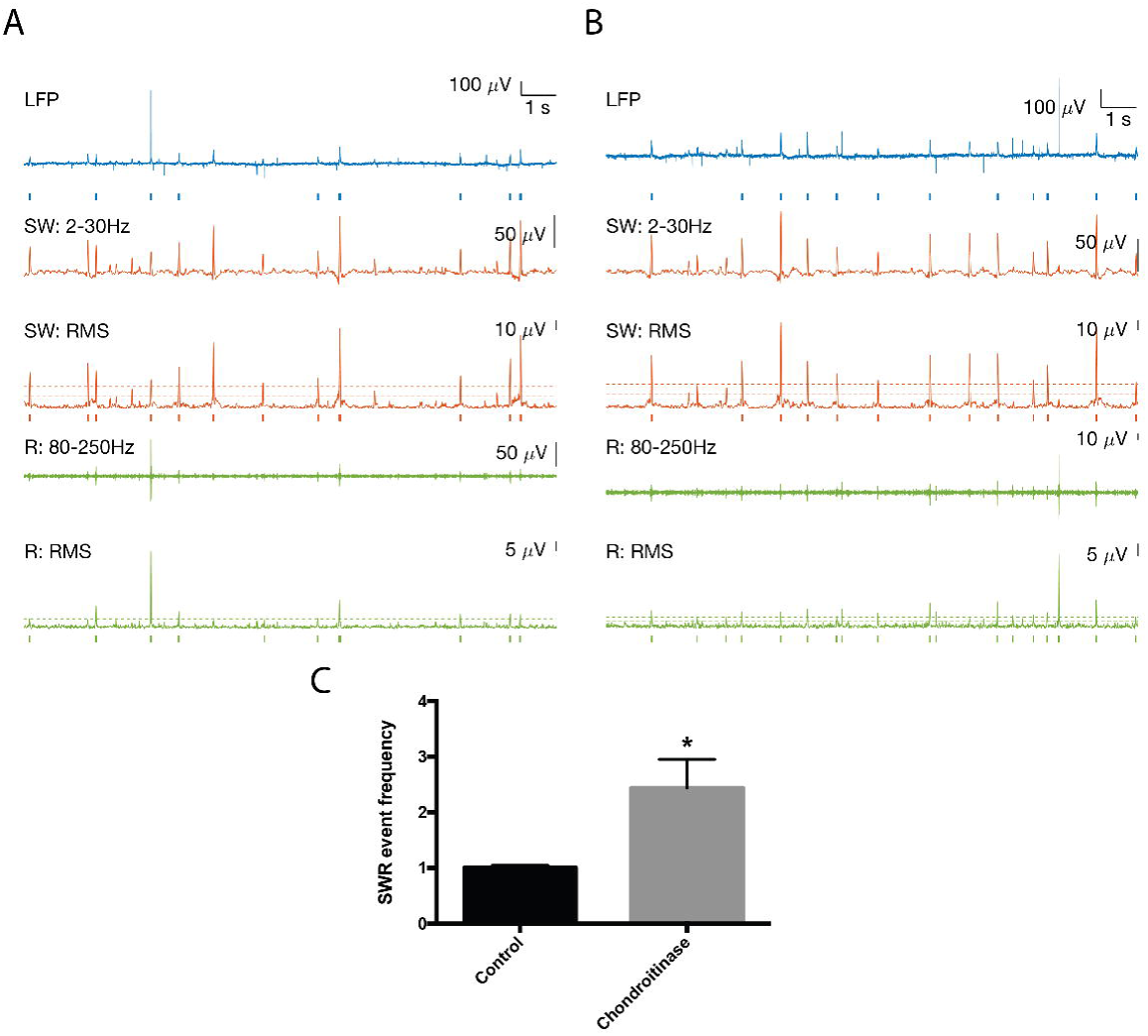
SWR event frequency in chondroitinase treated slices. Representative traces are shown in 4A and B, with SW events again in red and ripples in green. The fold change in average SWR event frequency results for 4 control and 4 treated slices is shown in 4C. The difference between control and treatment SWR frequency in 4C is significant (*p* < 0.05).

### V. Hypothetical schematic

In figure 5 we show a schematic representing a potential mechanism by which removal of perineuronal nets may increase SWR event frequency. In this model, an intact PNN can spatially restrict glutamatergic input to the PV neuron. This allows the PV neuron to inhibit activity of principal pyramidal cell. When PNN integrity is disrupted, however, the diffusion of glutamate and lateral mobility of post-synaptic receptors is likely increased, leading to reduced decreased excitatory input to PV neurons. This in turn leads to reduced inhibition, and thus increased excitability, of pyramidal cells.

**Figure 5.**
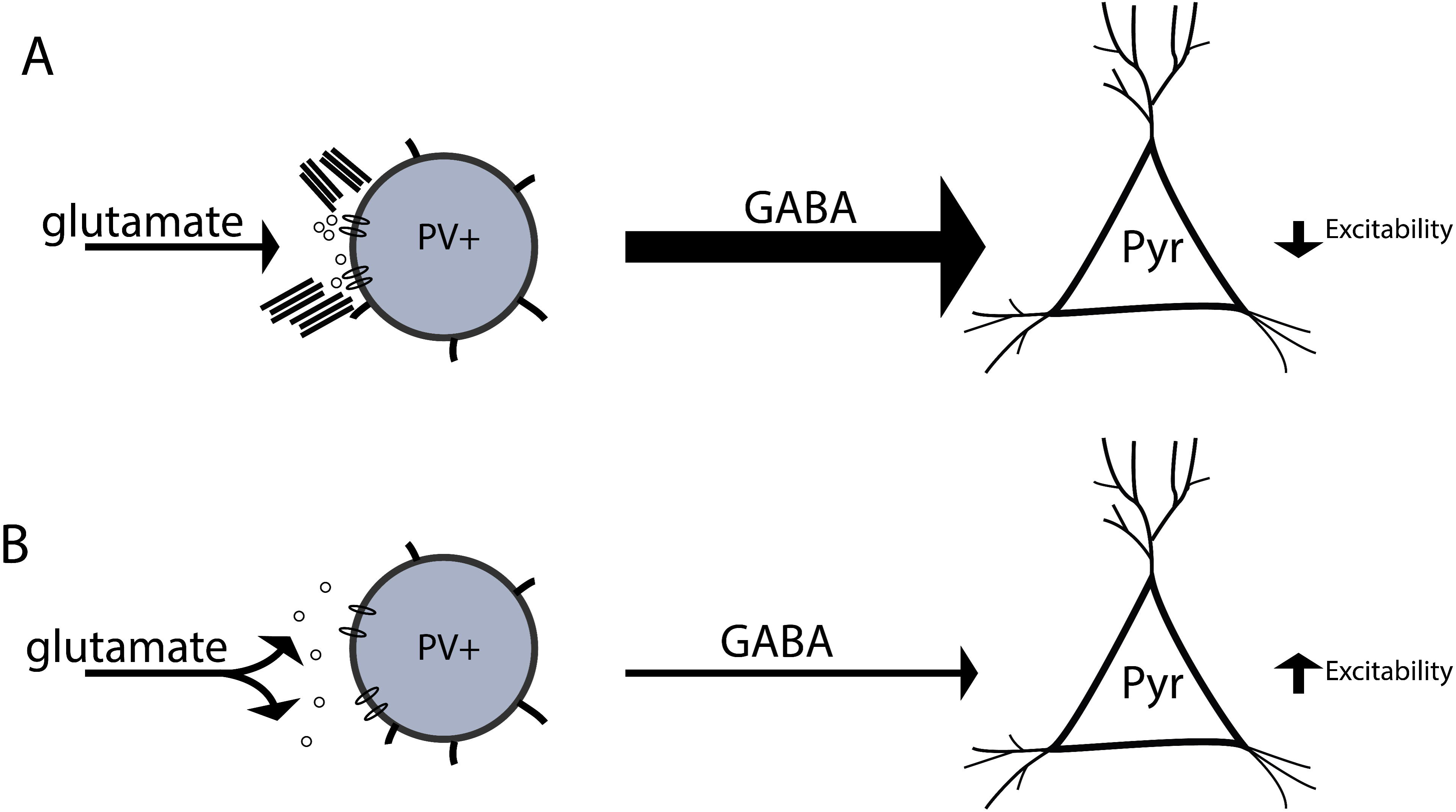
Hypothetical model by which PNN disruption influences the frequency of SWR events. In figure 6 we show a hypothetical model. A PV neuron with an intact PNN is shown at the top left. This neuron receives spatially localized glutamatergic input, allowing it to in turn inhibit pyramidal cell excitability. In contrast, a PV neuron with a disrupted PNN is shown at bottom left. In this scenario, glutamate may more easily diffuse from the synaptic cleft and/or glutamate receptors may demonstrate enhanced lateral mobility. A potential result is that this PV is less activated and thus less able to inhibit pyramidal cell excitability.

## Discussion

In the present study we observe that the frequency of SWR events is increased in hippocampal slices that are pretreated with proteases known to disrupt integrity of the PNN (Bikbaev et al., 2015). Given that increased pyramidal cell excitability is important to SWR initiation (Schlingloff et al., 2014), these data suggest that PNN integrity affects pyramidal cell excitability. In addition, since PV FSIs represent the predominant cell type known to be ensheathed by a PNN (Pantazopoulos and Berretta, 2016), these findings suggest that PNN digestion might indirectly affect pyramidal cell excitability through modulation of inhibitory input from the PV population. This possibility is supported by prior work with a mouse lacking GluAs on the PV population (Racz et al., 2009). This study demonstrated that when AMPA mediated excitation of PV interneurons was selectively suppressed, pyramidal cells were more likely to fire (Racz et al., 2009). The potential for PNN disruption to reduce PV mediated inhibition of pyramidal cells is supported by studies showing reduced excitability of PV cells, and a reduction in the frequency of inhibitory currents in pyramidal cells, following enzyme-mediated cleavage of PNNs (Slaker et al., 2015). A recent study also suggests loss of the PNN can decrease inhibition in the visual cortex (Lensjo et al., 2017). Of particular interest, this was associated with an increase in gamma frequency activity (Lensjo et al., 2017). Moreover, in related modeling work, it was demonstrated that reduced AMPA input to interneurons can lead to induction of SW bursts followed by high frequency gamma tails (Traub et al., 1996).

In terms of why disrupted PNN integrity would substantially reduce the ability of PV interneurons to inhibit pyramidal cell excitability, hypothetical mechanisms include the possibility that the PNN serves to properly localize glutamatergic input to this cell type (Bikbaev et al., 2015; Frischknecht et al., 2009). Glutamate receptors on PV interneurons are localized to soma and proximal dendrites, both of which are typically ensheathed by PNN. With PNN disruption released glutamate may more easily diffuse. In the absence of a robust PNN, receptor subunits might also be more motile and less precisely localized (Frischknecht et al., 2009). The negatively charged PNN may also buffer ions such as potassium (Bruckner et al., 1993; Hartig et al., 1999), and it has been proposed that removal of negative charge at the cell membrane could potentially affect the local electric field sensed by gating subunits of ion channels (Balmer, 2016).

Given that PV FSI activity is critical to ripple expression, it is of interest that despite the potential of PNN degradation to reduce PV FSI activity (Slaker et al., 2015), SWRs are well expressed in hyaluronidase and chondroitinase pre-treated slices. One possibility is that while PV cells are less excitable between SW events, allowing the same to occur more frequently, the PV population remains sufficiently excitable to express ripple activity when a SW initiating event occurs. Consistent with this, a prior study focused on PNN surrounded fast spiking neurons in the medial nucleus of the trapezoid body demonstrated that while PNN digestion reduced the gain of spike output in response to current steps in the background of Gaussian white noise currents, the ability of the neurons to fire up to 1000 Hz in response to current pulses was not affected (Balmer, 2016). In addition, work by Caputi *et al*. showed that reduced expression of GluA4 on FSIs was associated with increased firing of pyramidal cells but no alteration in the firing of FSIs during SWRs (Caputi et al., 2012).

Because PV interneurons represent the main cell type known to be ensheathed by a PNN, and because this cell type may be critical to SWR initiation and expression, our hypothetical mechanism (figure 5) is focused on changes in PV excitability. We acknowledge, however, that future studies will be necessary to examine the effect of PNN degradation on both PV cell and pyramidal cell activity during SWs and SWRs. The use of mice with ultrafast calcium indicators in specific cell populations could be particularly useful. These experiments will be important in that within CA2 hippocampus, a subpopulation of pyramidal cells is ensheathed by a PNN, and degradation of CA2 PNNs has been linked to recovery of the potential for long-term potentiation in this normally resistant region (Carstens et al., 2016). PNN degradation in this area might thus directly increase pyramidal cell activation. Recent work suggests that, as is the case for CA3, pyramidal cell activation within CA2 can initiate SWRs (Oliva et al., 2016).

In terms of the significance of PNN net integrity as an effector of SWR event frequency, expression and/or integrity of PNN components can be influenced by age, enrichment or disease (Berretta, 2012; Matuszko et al., 2017; Quattromani et al., 2017; Slaker et al., 2016a; Wiegand et al., 2016). Glial cells express critical components of the PNN, and expression of components is regulated by factors including neuronal activity (Howell and Gottschall, 2012; Matthews et al., 2002; McRae et al., 2007; Song and Dityatev, 2017). In addition, several PNN cleaving enzymes are expressed by neurons and glia (Brzdak et al., 2017; Deb and Gottschall, 1996; Dubey et al., 2017; Janusz et al., 2013; Pollock et al., 2014; Szklarczyk et al., 2002; Wiera et al., 2017; Yong et al., 1998). Soluble and transmembrane matrix metalloproteinases have been particularly well studied for their potential to disrupt PNN integrity, and their expression and/or activity is can be dramatically altered by physiological or pathological neuronal activity (Conant et al., 2010; Meighan et al., 2006; Nagy et al., 2006), disease states and select therapeutics (Szklarczyk et al., 2002). For example, MMP-9 levels are increased in response to seizure activity in varied models, and elevated MMP-9 has in turn been linked to reduced PNN integrity (Pollock et al., 2014). Though the physiology of epileptiform discharges and SWs differ in many respects, it is tempting to speculate that seizure induced PNN degradation may enhance pyramidal cell excitability important to both, and thus enhanced seizure risk as a function of poorly controlled epilepsy may follow from PNN disruption and previously described mossy fiber sprouting (Cronin et al., 1992; Golarai and Sutula, 1996). Disinhibition induced hyperexcitability and epileptiform like activity has been described in hippocampal cultures following treatment with hyaluronidase (Bikbaev et al., 2015; Vedunova et al., 2013). Moreover, in a study of post traumatic injury in rats, a loss of the PNN was associated with reduced inhibitory tone, leading the authors to suggest that PNN disruption might contribute to post traumatic epilepsy (Hsieh et al., 2016).

Of additional relevance are studies focused on psychiatric diseases including schizophrenia and major depression. These conditions have been associated with altered PNN integrity in combination with altered neuronal population activity. Major depression has been associated with both increased and decreased levels of PNN degrading enzymes, which may reflect the opposing influence of stress or inflammation in the former as opposed to reduced monoamine-stimulated protease expression in the latter. The antidepressant fluoxetine, which enhances monoaminergic tone, has been shown to increase MMP-9 levels and to reduce PNN integrity (Guirado et al., 2014). While multiple mechanisms likely contribute to fluoxetine’s antidepressant efficacy (Lee et al., 2015; Nestler et al., 1990; Nibuya et al., 1996), given that reduced excitatory/inhibitory balance has been implicated in the condition (Thompson et al., 2015), fluoxetine’s potential to reduce PNN integrity could contribute. Intriguingly, the fast acting antidepressant ketamine is thought to preferentially inhibit NMDA receptors on inhibitory interneurons (Quirk et al., 2009), suggesting that rapid and slow acting antidepressants might both target the activity of inhibitory interneurons albeit through temporally distinct mechanisms.

In summary, we demonstrate that the PNN disrupting enzymes hyaluronidase and chondroitinase can significantly increase the frequency of SWR events in horizontal hippocampal slices. We suggest that further study may be warranted to test the importance of PNNs on *in vivo* SWR dynamics and memory consolidation, in both normal physiology and disease states.

## Acknowledgements

We would like to acknowledge funding from the Von Matsch professorship for neurological diseases, the Dean’s Toulmin award, and the National Institutes of Health (NS083410). AC and PLB received support from T32 NS041231. ZS received support from a Chinese State Scholarship (201408220100).

## References

Alme CB, Miao C, Jezek K, Treves A, Moser EI, Moser MB. 2014. Place cells in the hippocampus: eleven maps for eleven rooms. Proc Natl Acad Sci U S A 111(52):18428–35.

Balmer TS. 2016. Perineuronal Nets Enhance the Excitability of Fast-Spiking Neurons. eNeuro 3(4).

Berretta S. 2012. Extracellular matrix abnormalities in schizophrenia. Neuropharmacology 62(3):1584–1597.

Bikbaev A, Frischknecht R, Heine M. 2015. Brain extracellular matrix retains connectivity in neuronal networks. Sci Rep 5:14527.

Bruckner G, Brauer K, Hartig W, Wolff JR, Rickmann MJ, Derouiche A, Delpech B, Girard N, Oertel WH, Reichenbach A. 1993. Perineuronal nets provide a polyanionic, glia-associated form of microenvironment around certain neurons in many parts of the rat brain. Glia 8(3):183–200.

Brzdak P, Wlodarczyk J, Mozrzymas JW, Wojtowicz T. 2017. Matrix Metalloprotease 3 Activity Supports Hippocampal EPSP-to-Spike Plasticity Following Patterned Neuronal Activity via the Regulation of NMDAR Function and Calcium Flux. Mol Neurobiol 54(1):804–816.

Buzsaki G. 1986. Hippocampal sharp waves: their origin and significance. Brain Res 398(2):242–52.

Buzsaki G. 2015. Hippocampal sharp wave-ripple: A cognitive biomarker for episodic memory and planning. Hippocampus 25(10):1073–188.

Caputi A, Fuchs EC, Allen K, Le Magueresse C, Monyer H. 2012. Selective Reduction of AMPA Currents onto Hippocampal Interneurons Impairs Network Oscillatory Activity. Plos One 7(6).

Carstens KE, Phillips ML, Pozzo-Miller L, Weinberg RJ, Dudek SM. 2016. Perineuronal Nets Suppress Plasticity of Excitatory Synapses on CA2 Pyramidal Neurons. J Neurosci 36(23):6312–20.

Ccedil;aliskan G, Schulz SB, Gruber D, Behr J, Heinemann U, Gerevich Z. 2015. Corticosterone and corticotropin-releasing factor acutely facilitate gamma oscillations in the hippocampus in vitro. European Journal of Neuroscience 41(1):31–44.

Colgin LL. 2016. Rhythms of the hippocampal network. Nat Rev Neurosci 17(4):239–49.

Conant K, Wang Y, Szklarczyk A, Dudak A, Mattson MP, Lim ST. 2010. Matrix metalloproteinase-dependent shedding of intercellular adhesion molecule-5 occurs with long-term potentiation. Neuroscience 166(2):508–21.

Cornez G, Madison FN, Van der Linden A, Cornil C, Yoder KM, Ball GF, Balthazart J. 2017. Perineuronal nets and vocal plasticity in songbirds: A proposed mechanism to explain the difference between closed-ended and open-ended learning. Dev Neurobiol.

Cronin J, Obenaus A, Houser CR, Dudek FE. 1992. Electrophysiology of dentate granule cells after kainate-induced synaptic reorganization of the mossy fibers. Brain Res 573(2):305–10.

Csicsvari J, Hirase H, Czurko A, Mamiya A, Buzsaki G. 1999. Fast network oscillations in the hippocampal CA1 region of the behaving rat. J Neurosci 19(16):RC20.

Deb S, Gottschall PE. 1996. Increased production of matrix metalloproteinases in enriched astrocyte and mixed hippocampal cultures treated with beta-amyloid peptides. J Neurochem 66(4):1641–7.

Dityatev A, Bruckner G, Dityateva G, Grosche J, Kleene R, Schachner M. 2007. Activity-dependent formation and functions of chondroitin sulfate-rich extracellular matrix of perineuronal nets. Dev Neurobiol 67(5):570–88.

Dragoi G, Tonegawa S. 2011. Preplay of future place cell sequences by hippocampal cellular assemblies. Nature 469(7330):397–401.

Dubey D, McRae PA, Rankin-Gee EK, Baranov E, Wandrey L, Rogers S, Porter BE. 2017. Increased metalloproteinase activity in the hippocampus following status epilepticus. Epilepsy Res 132:50–58.

Ego-Stengel V, Wilson MA. 2010. Disruption of ripple-associated hippocampal activity during rest impairs spatial learning in the rat. Hippocampus 20(1):1–10.

Eschenko O, Ramadan W, Molle M, Born J, Sara SJ. 2008. Sustained increase in hippocampal sharp-wave ripple activity during slow-wave sleep after learning. Learn Mem 15(4):222–8.

Filous AR, Tran A, Howell CJ, Busch SA, Evans TA, Stallcup WB, Kang SH, Bergles DE, Lee SI, Levine JM and others. 2014. Entrapment via synaptic-like connections between NG2 proteoglycan+ cells and dystrophic axons in the lesion plays a role in regeneration failure after spinal cord injury. J Neurosci 34(49):16369–84.

Franklin SL, Love S, Greene JR, Betmouni S. 2008. Loss of Perineuronal Net in ME7 Prion Disease.J Neuropathol Exp Neurol 67(3):189–99.

Frischknecht R, Heine M, Perrais D, Seidenbecher CI, Choquet D, Gundelfinger ED. 2009. Brain extracellular matrix affects AMPA receptor lateral mobility and shortterm synaptic plasticity. Nat Neurosci 12(7):897–904.

Gianfranceschi L, Siciliano R, Walls J, Morales B, Kirkwood A, Huang ZJ, Tonegawa S, Maffei L. 2003. Visual cortex is rescued from the effects of dark rearing by overexpression of BDNF. Proc Natl Acad Sci U S A 100(21):12486–91.

Girardeau G, Benchenane K, Wiener SI, Buzsaki G, Zugaro MB. 2009. Selective suppression of hippocampal ripples impairs spatial memory. Nat Neurosci 12(10):1222–3.

Golarai G, Sutula TP. 1996. Functional alterations in the dentate gyrus after induction of long-term potentiation, kindling, and mossy fiber sprouting. J Neurophysiol 75(1):343–53.

Guirado R, Perez-Rando M, Sanchez-Matarredona D, Castren E, Nacher J. 2014. Chronic fluoxetine treatment alters the structure, connectivity and plasticity of cortical interneurons. Int J Neuropsychopharmacol 17(10):1635–46.

Hartig W, Derouiche A, Welt K, Brauer K, Grosche J, Mader M, Reichenbach A, Bruckner G. 1999. Cortical neurons immunoreactive for the potassium channel Kv3.1b subunit are predominantly surrounded by perineuronal nets presumed as a buffering system for cations. Brain Res 842(1):15–29.

Hockfield S, Kalb RG, Zaremba S, Fryer H. 1990. Expression of neural proteoglycans correlates with the acquisition of mature neuronal properties in the mammalian brain. Cold Spring Harb Symp Quant Biol 55:505–14.

Howell MD, Gottschall PE. 2012. Lectican proteoglycans, their cleaving metalloproteinases, and plasticity in the central nervous system extracellular microenvironment. Neuroscience 217:6–18.

Hsieh TH, Lee HH, Hameed MQ, Pascual-Leone A, Hensch TK, Rotenberg A. 2016. Trajectory of Parvalbumin Cell Impairment and Loss of Cortical Inhibition in Traumatic Brain Injury. Cereb Cortex.

Janusz A, Milek J, Perycz M, Pacini L, Bagni C, Kaczmarek L, Dziembowska M. 2013. The Fragile X mental retardation protein regulates matrix metalloproteinase 9 mRNA at synapses. J Neurosci 33(46):18234–41.

Ji D, Wilson MA. 2007. Coordinated memory replay in the visual cortex and hippocampus during sleep. Nat Neurosci 10(1):100–7.

Koniaris E, Drimala P, Sotiriou E, Papatheodoropoulos C. 2011. Different effects of zolpidem and diazepam on hippocampal sharp wave-ripple activity in vitro. Neuroscience 175:224–34.

Kwok JC, Dick G, Wang D, Fawcett JW. 2011. Extracellular matrix and perineuronal nets in CNS repair. Dev Neurobiol 71(11):1073–89.

Lee SY, Chen SL, Chang YH, Chen PS, Huang SY, Tzeng NS, Wang CL, Wang LJ, Lee IH, Wang TY and others. 2015. Correlation of plasma brain-derived neurotrophic factor and metabolic profiles in drug-naive patients with bipolar II disorder after a twelve-week pharmacological intervention. Acta Psychiatr Scand 131(2):120–8.

Lensjo KK, Lepperod ME, Dick G, Hafting T, Fyhn M. 2017. Removal of Perineuronal Nets Unlocks Juvenile Plasticity Through Network Mechanisms of Decreased Inhibition and Increased Gamma Activity. J Neurosci 37(5):1269–1283.

Lin R, Kwok JC, Crespo D, Fawcett JW. 2008. Chondroitinase ABC has a long-lasting effect on chondroitin sulphate glycosaminoglycan content in the injured rat brain. J Neurochem 104(2):400–8.

Maier N, Nimmrich V, Draguhn A. 2003. Cellular and network mechanisms underlying spontaneous sharp wave-ripple complexes in mouse hippocampal slices. Journal of Physiology-London 550(3):873–887.

Matthews RT, Kelly GM, Zerillo CA, Gray G, Tiemeyer M, Hockfield S. 2002. Aggrecan glycoforms contribute to the molecular heterogeneity of perineuronal nets. J Neurosci 22(17):7536–47.

Matuszko G, Curreli S, Kaushik R, Becker A, Dityatev A. 2017. Extracellular matrix alterations in the ketamine model of schizophrenia. Neuroscience.

Maya Vetencourt JF, Sale A, Viegi A, Baroncelli L, De Pasquale R, O’Leary OF, Castren E, Maffei L. 2008. The antidepressant fluoxetine restores plasticity in the adult visual cortex. Science 320(5874):385–8.

Mayer J, Hamel MG, Gottschall PE. 2005. Evidence for proteolytic cleavage of brevican by the ADAMTSs in the dentate gyrus after excitotoxic lesion of the mouse entorhinal cortex. BMC Neurosci 6:52.

McRae PA, Rocco MM, Kelly G, Brumberg JC, Matthews RT. 2007. Sensory deprivation alters aggrecan and perineuronal net expression in the mouse barrel cortex. J Neurosci 27(20):5405–13.

Meighan SE, Meighan PC, Choudhury P, Davis CJ, Olson ML, Zornes PA, Wright JW, Harding JW. 2006. Effects of extracellular matrix-degrading proteases matrix metalloproteinases 3 and 9 on spatial learning and synaptic plasticity. J Neurochem 96(5):1227–41.

Mower GD. 1991. The effect of dark rearing on the time course of the critical period in cat visual cortex. Brain Res Dev Brain Res 58(2):151–8.

Nagy V, Bozdagi O, Matynia A, Balcerzyk M, Okulski P, Dzwonek J, Costa RM, Silva AJ, Kaczmarek L, Huntley GW. 2006. Matrix metalloproteinase-9 is required for hippocampal late-phase long-term potentiation and memory. J Neurosci 26(7):1923–34.

Nestler EJ, McMahon A, Sabban EL, Tallman JF, Duman RS. 1990. Chronic antidepressant administration decreases the expression of tyrosine hydroxylase in the rat locus coeruleus. Proc Natl Acad Sci U S A 87(19):7522–6.

Nibuya M, Nestler EJ, Duman RS. 1996. Chronic antidepressant administration increases the expression of cAMP response element binding protein (CREB) in rat hippocampus. J Neurosci 16(7):2365–72.

O’Keefe J, Dostrovsky J. 1971. The hippocampus as a spatial map. Preliminary evidence from unit activity in the freely-moving rat. Brain Res 34(1):171–5.

Oliva A, Fernandez-Ruiz A, Buzsaki G, Berenyi A. 2016. Role of Hippocampal CA2 Region in Triggering Sharp-Wave Ripples. Neuron 91(6):1342–55.

Oray S, Majewska A, Sur M. 2004. Dendritic spine dynamics are regulated by monocular deprivation and extracellular matrix degradation. Neuron 44(6):1021–30.

Pantazopoulos H, Berretta S. 2016. In Sickness and in Health: Perineuronal Nets and Synaptic Plasticity in Psychiatric Disorders. Neural Plast 2016:9847696.

Pantazopoulos H, Markota M, Jaquet F, Ghosh D, Wallin A, Santos A, Caterson B, Berretta S. 2015. Aggrecan and chondroitin-6-sulfate abnormalities in schizophrenia and bipolar disorder: a postmortem study on the amygdala. Transl Psychiatry 5:e496.

Papatheodoropoulos C, Sotiriou E, Kotzadimitriou D, Drimala P. 2007. At clinically relevant concentrations the anaesthetic/amnesic thiopental but not the anticonvulsant phenobarbital interferes with hippocampal sharp wave-ripple complexes. BMC Neurosci 8:60.

Pizzorusso T, Medini P, Berardi N, Chierzi S, Fawcett JW, Maffei L. 2002. Reactivation of ocular dominance plasticity in the adult visual cortex. Science 298(5596):1248–51.

Place R, Farovik A, Brockmann M, Eichenbaum H. 2016. Bidirectional prefrontal-hippocampal interactions support context-guided memory. Nat Neurosci 19(8):992–4.

Pollock E, Everest M, Brown A, Poulter MO. 2014. Metalloproteinase inhibition prevents inhibitory synapse reorganization and seizure genesis. Neurobiol Dis 70:21–31.

Quattromani MJ, Pruvost M, Guerreiro C, Backlund F, Englund E, Aspberg A, Jaworski T, Hakon J, Ruscher K, Kaczmarek L and others. 2017. Extracellular Matrix Modulation Is Driven by Experience-Dependent Plasticity During Stroke Recovery. Mol Neurobiol.

Quirk MC, Sosulski DL, Feierstein CE, Uchida N, Mainen ZF. 2009. A defined network of fast-spiking interneurons in orbitofrontal cortex: responses to behavioral contingencies and ketamine administration. Front Syst Neurosci 3:13.

Racz A, Ponomarenko AA, Fuchs EC, Monyer H. 2009. Augmented hippocampal ripple oscillations in mice with reduced fast excitation onto parvalbumin-positive cells. J Neurosci 29(8):2563–8.

Rothschild G, Eban E, Frank LM. 2017. A cortical-hippocampal-cortical loop of information processing during memory consolidation. Nat Neurosci 20(2):251–259.

Sadowski JH, Jones MW, Mellor JR. 2011. Ripples make waves: binding structured activity and plasticity in hippocampal networks. Neural Plast 2011:960389.

Schlingloff D, Kali S, Freund TF, Hajos N, Gulyas AI. 2014. Mechanisms of sharp wave initiation and ripple generation. J Neurosci 34(34):11385–98.

Siapas AG, Wilson MA. 1998. Coordinated interactions between hippocampal ripples and cortical spindles during slow-wave sleep. Neuron 21(5):1123–8.

Skaggs WE, McNaughton BL, Wilson MA, Barnes CA. 1996. Theta phase precession in hippocampal neuronal populations and the compression of temporal sequences. Hippocampus 6(2):149–72.

Slaker M, Barnes J, Sorg BA, Grimm JW. 2016a. Impact of Environmental Enrichment on Perineuronal Nets in the Prefrontal Cortex following Early and Late Abstinence from Sucrose Self-Administration in Rats. Plos One 11(12).

Slaker M, Churchill L, Todd RP, Blacktop JM, Zuloaga DG, Raber J, Darling RA, Brown TE, Sorg BA. 2015. Removal of perineuronal nets in the medial prefrontal cortex impairs the acquisition and reconsolidation of a cocaine-induced conditioned place preference memory. J Neurosci 35(10):4190–202.

Slaker ML, Harkness JH, Sorg BA. 2016b. A standardized and automated method of perineuronal net analysis using Wisteria floribunda agglutinin staining intensity. IBRO Reports 1:54–60.

Song I, Dityatev A. 2017. Crosstalk between glia, extracellular matrix and neurons. Brain Res Bull.

Sorg BA, Berretta S, Blacktop JM, Fawcett JW, Kitagawa H, Kwok JC, Miquel M. 2016. Casting a Wide Net: Role of Perineuronal Nets in Neural Plasticity. J Neurosci 36(45):11459–11468.

Szklarczyk A, Lapinska J, Rylski M, McKay RD, Kaczmarek L. 2002. Matrix metalloproteinase-9 undergoes expression and activation during dendritic remodeling in adult hippocampus. J Neurosci 22(3):920–30.

Thompson SM, Kallarackal AJ, Kvarta MD, Van Dyke AM, LeGates TA, Cai X. 2015. An excitatory synapse hypothesis of depression. Trends Neurosci 38(5):279–94.

Traub RD, Whittington MA, Colling SB, Buzsaki G, Jefferys JG. 1996. Analysis of gamma rhythms in the rat hippocampus in vitro and in vivo. J Physiol 493 ( Pt 2):471–84.

Vedunova M, Sakharnova T, Mitroshina E, Perminova M, Pimashkin A, Zakharov Y, Dityatev A, Mukhina I. 2013. Seizure-like activity in hyaluronidase-treated dissociated hippocampal cultures. Front Cell Neurosci 7:149.

Whittington MA, Traub RD, Jefferys JG. 1995. Synchronized oscillations in interneuron networks driven by metabotropic glutamate receptor activation. Nature 373(6515):612–5.

Wiegand JPL, Gray DT, Schimanski LA, Lipa P, Barnes CA, Cowen SL. 2016. Age Is Associated with Reduced Sharp-Wave Ripple Frequency and Altered Patterns of Neuronal Variability. Journal of Neuroscience 36(20):5650–5660.

Wiera G, Nowak D, van Hove I, Dziegiel P, Moons L, Mozrzymas JW. 2017. Mechanisms of NMDA Receptor- and Voltage-Gated L-Type Calcium Channel-Dependent Hippocampal LTP Critically Rely on Proteolysis That Is Mediated by Distinct Metalloproteinases. J Neurosci 37(5):1240–1256.

Wilson MA, McNaughton BL. 1994. Reactivation of hippocampal ensemble memories during sleep. Science 265(5172):676–9.

Yong VW, Krekoski CA, Forsyth PA, Bell R, Edwards DR. 1998. Matrix metalloproteinases and diseases of the CNS. Trends Neurosci 21(2):75–80.

